# Dysregulated acetylcholine-mediated dopamine neurotransmission in the eIF4E Tg mouse model of autism spectrum disorders

**DOI:** 10.1101/2024.01.29.577831

**Authors:** Josep Carbonell-Roig, Alina Aaltonen, Veronica Cartocci, Avery McGuirt, Eugene Mosharov, Jan Kehr, Ori J. Lieberman, David Sulzer, Anders Borgkvist, Emanuela Santini

## Abstract

Autism Spectrum Disorders (ASD) consist of diverse neurodevelopmental conditions where core behavioral symptoms are critical for diagnosis. Altered dopamine neurotransmission in the striatum has been suggested to contribute to the behavioral features of ASD. Here, we examine dopamine neurotransmission in a mouse model of ASD characterized by elevated expression of the eukaryotic initiation factor 4E (eIF4E), a key regulator of cap-dependent translation, using a comprehensive approach that encompasses genetics, behavior, synaptic physiology, and imaging. The results indicate that increased eIF4E expression leads to behavioral inflexibility and impaired striatal dopamine release. The loss of normal dopamine neurotransmission is due to a defective nicotinic receptor signaling that regulates calcium dynamics in dopaminergic axons. These findings reveal an intricate interplay between eIF4E, DA neurotransmission, and behavioral flexibility, provide a mechanistic understanding of ASD symptoms and offer a foundation for targeted therapeutic interventions.

## Introduction

Autism Spectrum Disorders (ASD) encompass a diverse set of polygenic neurodevelopmental conditions defined by core behavioral symptoms, such as repetitive, stereotyped behaviors and abnormal social interactions^1^. The behavioral manifestations within ASD are heterogeneous and are often complicated by comorbidities. Thus, the clinical diagnosis of ASD is challenging since it primarily relies on the identification of this complex behavioral symptomatology^2^.

Emerging clinical and preclinical research aimed at determining brain alterations associated with ASD symptomatology suggests that the behavioral manifestations of ASD may stem from altered synaptic functions in the striatum: the main receiving nucleus of the basal ganglia^3-5^. Dopamine (DA) neurotransmission is crucial for striatal physiology as it plays a key role in selecting and transmitting motor patterns, and contributes to reward-based learning, motivation, and salience. Disrupted DA signaling has been linked to hyperactivity, inattention, and stereotyped behaviors found in several neurodevelopmental disorders^6-8^. Notably, abnormal DA signaling is observed in ASD patients and animal models^6-9^, though the precise mechanisms linking altered striatal DA release and ASD symptomatology remain the subject of ongoing research.

Syndromic^10-14^ and idiopathic^15-18^ forms of ASD have been linked to dysregulated mammalian target of rapamycin (mTOR) signaling. The mTOR pathway regulates essential biochemical processes, including activity-dependent protein synthesis by releasing the eIF4E-mRNAs complexes from the inhibitory eIF4E-binding protein 2 (4E-BP2)^19,20^. eIF4E binds the methyl-7-guanosine (m^7^G) cap on the 5’ end of eukaryotic mRNAs and is essential for physiological cap-dependent translation^19,21^. In neurons, eIF4E-dependent translation is critical for synaptic and structural plasticity, processes that are disrupted in preclinical models and individuals with ASD^10,22,23^.

Consistent with the evidence that altered mTOR signaling and cap-dependent translation are pathogenic molecular alterations shared by ASD, mutations in the *EIF4E* gene, which encodes eIF4E, have been linked to non-syndromic ASD^24^. Genetic variations on chromosome 4q, which contains the *EIF4E* gene, have been described in ASD patients^25,26^. Notably, recent findings have revealed elevated levels of eIF4E-dependent translation in postmortem brains of individuals with ASD^14^.

The central role of eIF4E-dependent translation in the synaptic, structural, and behavioral impairments that are characteristic of ASD has been demonstrated with preclinical models. Mice with global (referred to as eIF4E Tg)^27^ or microglia-specific^28^ overexpression of eIF4E as well as constitutive genetic deletion of 4E-BP2^29^, showed that aberrant brain eIF4E-dependent translation is associated with ASD-like behaviors, synaptic and structural abnormalities consistent with ASD pathology. Moreover, in these models, causality was established through brain infusions of 4E-GI, an eIF4E inhibitor that reduces cap-dependent translation, restored synaptic function and normalized ASD-behaviors in the eIF4E Tg and 4E-BP2 KO mice^27,29^.

Here, we took advantage of the eIF4E Tg mice^27^ to test the hypothesis that behavioral inflexibility is linked to defective DA neurotransmission. Our research employed a combination of genetic, synaptic physiology and imaging methods to reveal that increased eIF4E expression results in reduced DA release in the striatum. Furthermore, our investigation identified a mechanism reliant on impaired nicotinic receptors (nAChR) and axonal Ca^2+^ dynamics. Here, we provide strong evidence for altered DA neurotransmission as a pathological striatal mechanism in ASD symptomatology.

## Results

### The eIF4E Tg mice exhibit aberrant reversal learning

Striatal DA is implicated in cognitive flexibility^30-32^, a feature commonly compromised in ASD patients^30,33-35^. Our previous studies have shown that the eIF4E Tg mice exhibit impaired reversal learning and cognitive flexibility in behavioral tasks based on negative reinforcement, for instance avoiding a punishment such as cold water or foot shock^27^. However, it remains unknown whether the behavioral inflexibility of the eIF4E Tg mice extends to their ability to adapt to novel contingencies for the purpose of achieving positive outcomes, such as rewards.

To address this question, we employed the four-choice odor discrimination task, a reversal learning paradigm based on a naturalistic reward (e.g., food), which is sensitive to changes in DA neurotransmission^36,37^. In this task, the mice initially learn to associate a food reward with a specific odor (O1-food in the discrimination phase; Figure 1a). The learned association is then challenged in the subsequent reversal phase, where the food is paired with a different odor (O2-food in the reversal phase; Figure 1a).

**Figure 1.**
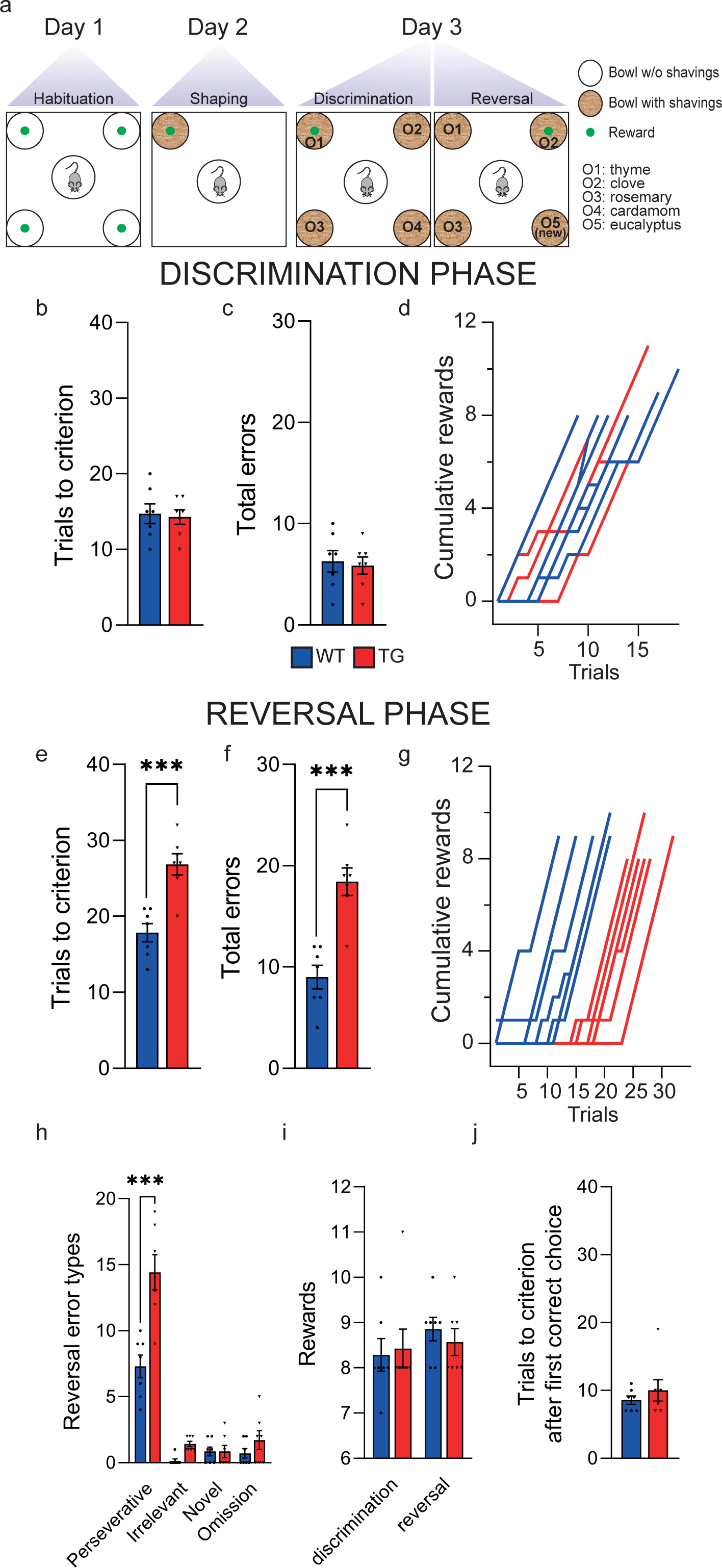
The eIF4E Tg mice display cognitive inflexibility in the four-choice odor discrimination task. (**a**) Schematic of the four-choice odor discrimination task. (**b**-**c**) Acquisition learning of O1-food association during discrimination, presented as number of (**b**) trials to reach criterion (eight out of ten), (**c**) total errors and (**d**) cumulative rewards plot. Unpaired, two-tailed *t-*test for (**b**) t_12_=0.3, ns and (**c**) t_12_=0.3, ns. (**e**-**g**) Reversal learning of O2-food association during test reversal, presented as number of (**e**) trials to reach criterion (eight out of ten), (**f**) total errors and (**g**) cumulative rewards plot. Unpaired, two-tailed *t-*test for (**e**) t_12_=4.9, ***p<0.001 and (**f**) t_12_=5.3, ***p<0.001. (**h**) Error types during task reversal determined by the choice to dig in a specific odor containing pot and classified as follows: perseverative (odor rewarded during discrimination, i.e., O1), irrelevant (odor that was never rewarded), novel (new introduced odor, never rewarded, i.e., O5,), and omission (no choice within 3 minutes). Two-way RM ANOVA, Genotype: F_(1, 12)_=27.9, p<0.001; Error Types: F_(1,24, 14,86)_=107.8, p<0.0001; interaction Genotype:Error types: F_(3, 36)_=11.5, p<0.0001; ***p<0.001, Bonferroni’s multiple comparison test. (**i**) Total number of food rewards obtained during the task discrimination and reversal. Unpaired, two-tailed *t-* test, t_12_=0.3, ns and t_12_=0.7, ns, respectively. (**j**) Number of trials to reach criterion after the first correct choice. Unpaired, two-tailed *t-*test for discrimination, t_12_=0.8, ns. For all graphs, bars represent group averages expressed as mean ± SEM and dots represent values for individual mice. Wildtype (WT) and eIF4E Tg mice (TG) are depicted by blue and red bars, respectively. N= 7 mice/ genotype. ns= not significant.

The eIF4E Tg mice displayed intact acquisition learning as evidenced by a similar number of trials required to reach criterion (eight successful food retrievals out of ten; Figure 1b) and a similar number of errors (Figure 1c) compared to their wildtype littermates. Moreover, plots illustrating the cumulative rewards obtained by the mice over the different trails indicated that the curves for both wildtype and eIF4E Tg mice overlapped (Figure 1d). In contrast, during the reversal phase, when the food was associated with a different odor (O2-food), the eIF4E Tg mice required a significantly higher number of trials to reach criterion (Figure 1e), made more errors (Figure 1f) and display a distinct, right-shifted non-overlapping cumulative reward curve (Figure 1g) compared to the wildtype mice. Notably, perseverative errors were the most frequent error type for all mice in the reversal phase of the test, but the eIF4E Tg mice exhibited significantly more perseverative errors than their wildtype littermates (Fig 1h). These results indicate that eIF4E Tg mice have impaired behavioral adaptation in response to altered environmental conditions.

To determine if hunger status impacted the performance of the eIF4E Tg mice in the four-choice odor discrimination task, we examined the total number of food rewards obtained during the task and the trials needed to reach criterion after their first correct choice. The results showed no significant differences between the eIF4E Tg and wildtype controls mice in both discrimination and reversal phases regarding the total food rewards (Figure 1i) and the trials to meet criterion after the first correct choice (Figure 1j). Overall, these findings suggest that hunger status is an unlikely factor contributing to the behavioral inflexibility of the eIF4E Tg mice.

In summary, the eIF4E Tg mice show inflexible behavior, as evidenced by their increased trials needed to adapt a learned behavior in response to changes in the environment.

### Reduced DA release in the dorsal striatum of eIF4E Tg mice

The eIF4E Tg mice exhibited altered reversal learning in the four-choice odor discrimination task. Similar results were previously shown to be dependent on altered DA release, excitation-inhibition (E/I) balance, firing pattern and morphology of DA neurons^36,37^. Therefore, we hypothesized that eIF4E Tg mice exhibit deficits in dopaminergic function. We characterized sub-second DA release using Fast-scan cyclic voltammetry (FSCV) following local electrical stimulation of acute striatal slices from the eIF4E Tg mice and their wild-type littermates. Beside single pulse stimulation, we also employed train stimulation (5 pulses at 40Hz; 5p40Hz) which emulates phasic firing of DA neurons and falls within the range that maximizes striatal DA release^38^.

Both single pulse and train stimulation revealed a significant reduction in the peak amplitude of DA efflux in the dorsal striatum of eIF4E Tg mice (Figure 2a and b).

**Figure 2.**
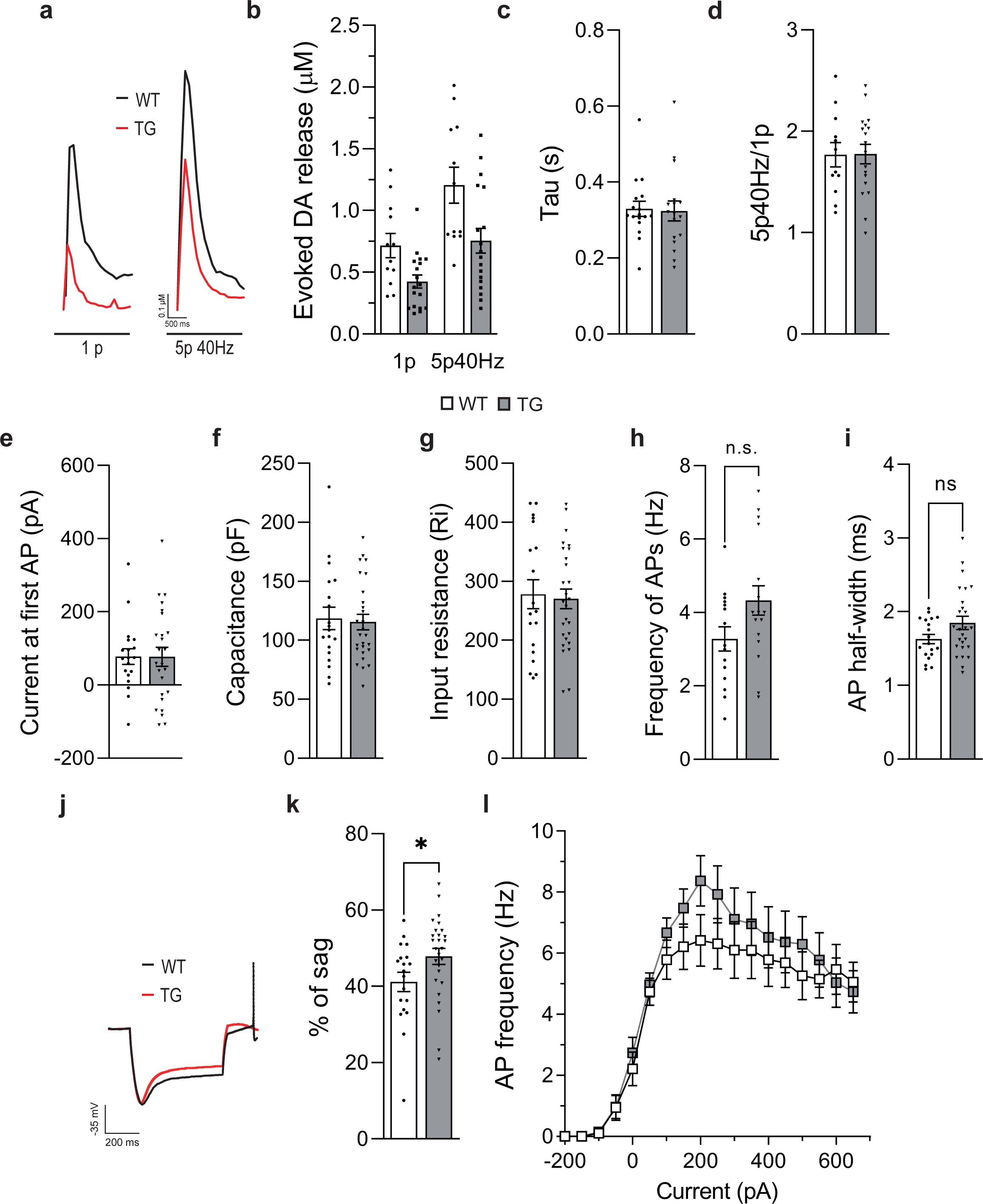
Altered striatal DA release but intact intrinsic properties and excitability of midbrain DA neurons of the eIF4E Tg mice. (**a-d**) FSCV recordings in the dorsal striatum of acute corticostriatal slices. (**a**) Representative traces of FSCV recordings presented in (b); wildtype (WT, black) and eIF4E (TG, red). (**b**) Peak concentrations of evoked DA release following a single pulse (1p) or 5 pulses at 40 Hz (5p40Hz). Two-way RM ANOVA, Genotype: F_(1, 28)_=7.85, p<0.01; Stimulation: F_(1, 28)_=72.9, p<0.0001; interaction Genotype:Stimulation: F_(1, 28)_=2.76, p=0.6, ns. (**c**) Time constant (Tau) of the exponential decay of the DA transients evoked by single pulse stimulus. Unpaired, two-tailed *t-*test, t_28_=0.5, ns. (**d**) Ratio of the peak DA release evoked by 5p40Hz over 1p, referring to (b). Unpaired, two-tailed *t-*test, t_28_=0.04, ns. N= 12-18 slices/ 3-4 mice/ genotype. (**e-g, i** and **j-l**) Whole-cell current-clamp and (**h**) cell-attached recordings of DA neurons in acute midbrain slices. (**e**) Current at the first action potential (AP). Unpaired, two-tailed *t-*test, t_42_=0.016, ns. (**f**) Capacitance. Unpaired, two-tailed *t-*test, t_44_=0.27, ns. (**g**) Input resistance. Unpaired, two-tailed *t-*test, t_44_=0.27, ns. (**h**) Number of AP in 10 seconds (Frequency). Unpaired, two-tailed *t-*test, t_30_=1.99, ns. (**i**) AP half-width. Unpaired, two-tailed *t-*test, t_43_=1.86, ns. (**j**) Representative voltage responses to -300pA injected current; wildtype (WT, black) and eIF4E (TG, red). (**k**) Ratio of the instantaneous versus steady state voltage response to -300pA current injection (% of sag). Unpaired, two-tailed *t-*test, t_44_=2.1, *p<0.05. (**l**) Rate of AP (Hz) in response to a series of current steps of increasing amplitude. Two-way RM ANOVA, Genotype: F_(1, 44)_=0.6, p=0.5, ns; Current steps: F_(2,65 116,6)_=66.2, p<0.0001; interaction Genotype:Current steps: F_(19, 836)_=0.8, P=0.5 ns. N= 15-27 neurons/ 3 mice/ genotype. For all graphs, bars represent group averages expressed as mean ± SEM and dots represent values for individual slices (**a**-**d**) or neurons (**e**-**l**). Wildtype (WT) and eIF4E Tg mice (TG) are depicted by white and grey bars, respectively. ns= not significant.

The observed decrease in DA amplitude was not a result of increased reuptake, as the decay constant (tau) of the DA peak, which is influenced by the efficacy of the dopamine reuptake transporter (DAT)^39^, was similar in both wild-type and eIF4E Tg mice (Figure 2c).

We found that DA efflux is dynamically regulated by increasing the stimulation paradigm in both genotypes as indicated by the similar train/single pulse ratio (Figure 2d). Since high frequency stimulation increases DA release by enhancing net axonal Ca^2+^ influx through voltage-gated Ca2+ channels^38,40^, our results indicate that there is no alteration in activity-dependent DA release related to major deficits in Ca^2+^ influx dynamics.

To examine if the reduced DA release in eIF4E Tg depends on altered regulation of release by D2 autoreceptors (D2R), which provide presynaptic feedback inhibition of axonal DA release^41-44^, we applied sulpiride (2 µM), a D2R-like antagonist to the slices while recording stimulus-evoked DA release. Notably, sulpiride did not have an impact on DA release in response to single-pulse stimulation in either genotype (Figure S1a), indicating that the reduced DA release in eIF4E Tg mice is not dependent on altered D2-autoreceptor activity.

Altogether, these results suggest that the reduced evoked DA release in the striatum of the eIF4E Tg mice is not attributable to a defective DAT and D2R function in the DA axonal terminals.

### Unaltered DA metabolism and electrophysiological properties of midbrain DA neurons in the eIF4E Tg mice

Since the availability of DA for release is also regulated by the synthesis, degradation, and vesicular storage of DA within axonal terminals^45^, our initial focus was to determine if alterations in these biochemical processes contributed to deficits in DA release observed in the eIF4E Tg mice. We employed quantitative Western blotting to measure key proteins involved in DA synthesis (i.e., tyrosine hydroxylase (TH)), vesicular loading (i.e., vesicular monoamine transporter 2 (vMAT2)), and reuptake (i.e., DAT) in the striatum^46,47^. Our analysis revealed that the levels of all the analyzed proteins were consistent between the eIF4E Tg mice and their wildtype littermates (Figure S1b and c). Immunostaining for TH also revealed a comparable density of midbrain DA neurons between genotypes (Figure S1e and f). Moreover, liquid chromatography tandem mass spectrometry (UHPLC-MS/MS) further established that the striatal levels of DA and its acidic metabolites 3,4-dihydroxyphenylacetic acid (DOPAC) and homovanillic acid (HVA) remained unaltered in the eIF4E Tg mice (Figure S1d).

While DA release is robustly controlled at the axonal level, the activity of the cell bodies is crucial for the overall regulation of DA neurotransmission^45,48-52^. Therefore, we extended our investigation to determine whether overexpression of eIF4E alters the physiology of midbrain DA neurons.

Cell-attached and whole-cell patch clamp recordings of DA neurons in acute midbrain slices^53,54^ demonstrated no significant differences in membrane properties or tonic activity of DA neurons (Figure 2e-i), aside from a modest increase in a voltage sag in response to hyperpolarizing currents (Figure 2j and k) suggesting a stronger activation of HCN channels^53-55^. Finally, action potential frequency evoked by current injection (IF curves) was not different between genotypes, indicating that DA neuron excitability remained unchanged in eIF4E Tg mice (Figure 2l).

Altogether, these findings demonstrate that the eIF4E Tg mice exhibited reduced striatal DA release without significant DA neuron pathophysiology.

### Impaired DA release triggered by acetylcholine (ACh) interneurons in the eIF4E Tg mice

A growing body of evidence suggests that striatal DA release is modulated by local heterosynaptic regulatory mechanisms at the DA terminals^38,45,56,57^. Recently it has been established that synchronous activation of ACh interneurons and the subsequent ACh release can trigger heterosynaptic DA release through the activation of β2 nicotinic receptors (β2-nAChR) on DA axons^58-61^. Thus, we examined if the reduced DA release in the eIF4E Tg mice originates from changes in heterosynaptic control of DA efflux by ACh.

First, we performed FSCV in the presence of the β2-nAChR antagonist dihydro-β-erythroidine (DHβE, 1 µM), which has been shown to significantly reduce electrically evoked DA release^58-61^. As predicted, bath application of DHβE reduced DA release triggered by single pulse stimulus in acute striatal slices obtained from wildtype mice (Figure 3a and b). In the eIF4E Tg mice the effect of DHβE was attenuated (Figure 3a and b), indicating a reduced effect of ACh on DA release (DHβE-sensitive DA release amounted to WT=0.38±0.05 µM and TG=0.18±0.04 µM; unpaired, two-tailed *t-*test, t_18_=2.94, **p<0.01).

**Figure 3.**
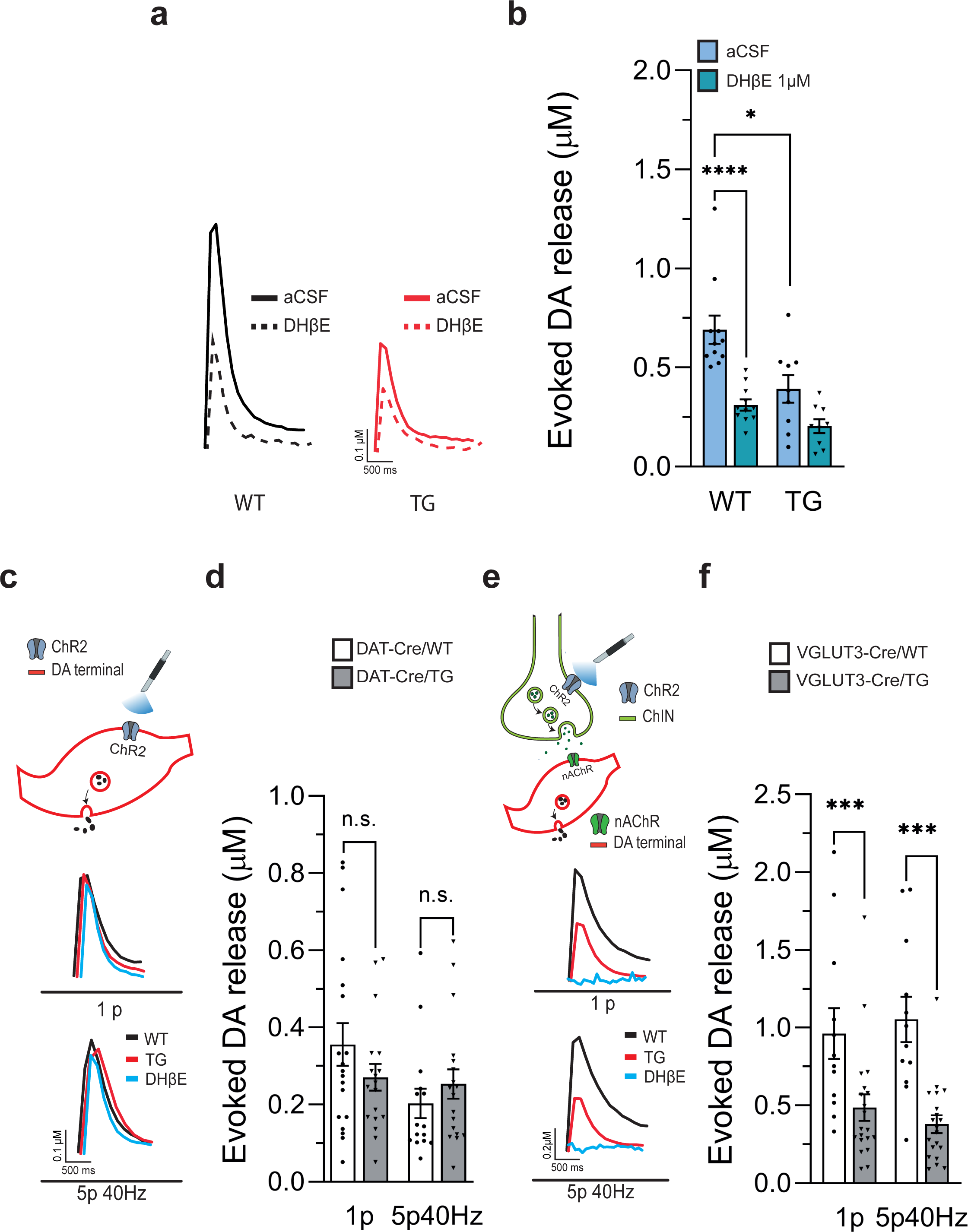
Impaired DA release induced by ACh in the eIF4E Tg mice. (**a-b**) FSCV recordings with DHβE (1µM), an inhibitor of β2 subunit-containing nAChR (β2-nAChR), in the dorsal striatum of acute corticostriatal slices (**a**) Representative traces of FSCV recordings presented in (b); wildtype (WT, black) and eIF4E Tg (TG, red) before (aCSF, continuous lines) and after (DHβE, dotted lines) treatment. (**b**) Peak concentrations of evoked DA release in the presence of DHβE (dark blue) or vehicle (aCSF; light blue) following a single pulse stimulus. Two-way RM ANOVA, Genotype: F_(1, 18)_=8.1, p=0.01; Treatment: F_(1,18)_=75.5, p<0.0001; interaction Genotype:Treatment: F_(1, 18)_=8.6, p=0,009; ***p<0.001, ****p<0.0001 and *p<0.05, Multiple comparison’s post-hoc test. N= 7-9 slices/ 3-4 mice/ genotype. (**c-e**) Selective optogenetic stimulation of DA axons or (**f-h**) striatal ACh interneurons and FSCV recordings in the dorsal striatum of acute corticostriatal slices. (**c and f**) Schematic of the experiments and (**d** and **g**) representative traces of FSCV recordings presented in (**e**) and (**h)**, respectively; wildtype (WT, black), eIF4E Tg (TG, red), DHβE (blue). DHβE traces included to illustrate the dependency of DA release on ACh-nAChR specifically in (**f-h**). Peak concentrations of evoked DA release following a single pulse (1p) or 5 pulses at 40 Hz (5p40Hz) of blue light stimulating channelrhodopsin 2 (ChR2) expressed in DA terminals (**e**) or ACh interneuron (**h**). Two-way RM ANOVA for (**e**) Genotype: F_(1, 66)_=0.2, p=0.7 ns; Stimulation: F_(1, 66)_=3.8, p=0.06 ns; interaction Genotype:Stimulation: F_(1, 66)_=2.5, p=0.1 ns. N= 19-18 slices/ 4 mice/ genotype; (**h**) Genotype: F_(1, 30)_=15.2, p=0.0005; Stimulation: F_(1, 30)_=0.05, p=0.8 ns; interaction Genotype:Stimulation: F_(1, 30)_=8.4, p=0.007; ***p<0.001, Multiple comparison’s post-hoc test. N= 12-20 slices/ 4-5 mice/ genotype. For all graphs, bars represent group averages expressed as mean ± SEM and dots represent values for individual slices. ns= not significant.

To isolate the direct impact of altered DA release induced by ACh in eIF4E Tg mice, we selectively expressed the light-activated ion channel channelrhodopsin 2 (ChR2) in DA terminals (Figure 3c, d and e) or in ACh interneurons (Figure 3f, g and h). For the selective targeting of ChR2 in DA terminals, we bilaterally injected an AAV5 carrying Cre-inducible ChR2 into the substantia nigra of mice generated by crossing eIF4E Tg with DAT-*Cre* animals (DAT-*Cre*/WT and -/TG; see materials and methods)^62,63^. The selective expression of ChR2 in DA axons was confirmed by colocalization of mCherry expressed by AAV5-ChR2 with TH via immunohistochemistry (Figure S2a).

For specific targeting of ChR2 in striatal ACh interneurons, we took advantage of their selective expression of VGLUT3^64,65^. Accordingly, an AAV5 with *Cre*-inducible ChR2 was bilaterally injected into the striatum of eIF4E Tg and wildtype mice expressing *Cre* in VGLUT3-positive ACh interneurons (VGLUT3-Cre/WT and - /TG, respectively; see materials and methods)^66^. Selective expression of *Cre* in striatal ACh interneurons was confirmed by stereotaxic striatal injection of AAV5- DIO-YFP and immunostaining, revealing near-complete colocalization of the *Cre*-driven YFP with choline acetyltransferase (ChAT; Figure S2b and c), a key enzyme for acetylcholine synthesis in ACh interneurons^64^.

Next, we prepared acute striatal slices from DAT-*Cre*/WT and -/TG mice specifically expressing ChR2 in DA neurons. In this preparation, light stimulation selectively activates ChR2 on the axons of DA neurons, leading to transient DA release detected by FSCV. Our findings indicate that optogenetically-induced DA release evoked by single-pulse or train stimulation was not impaired in mice overexpressing eIF4E (Figure 3d and e).

In contrast, in striatal slices from mice expressing ChR2 in VGLUT3+ ACh interneurons, light-evoked DA release requires the selective activation of ACh interneurons and functional β2-nAChR on DA axons^58,61^. Indeed, light-evoked DA release is completely abolished by the bath application of DHβE (1 µM; Figure 3g). Importantly, we observed impaired light-evoked DA release triggered by ACh in mice overexpressing eIF4E, induced either by a single pulse or train (5p40Hz) stimulation (Figure 3g and h). These results confirm that the intrinsic mechanisms regulating axonal DA release in the eIF4E Tg mice are intact; the observed deficit in these mice is specifically attributed to compromised axonal DA release triggered by acetylcholine.

### Unaltered firing properties, release, and total striatal content of ACh in the eIF4E Tg mice

To investigate whether the reduction in DA release triggered by ACh is linked to functional deficiencies in ACh interneurons of the eIF4E Tg mice, we conducted cell-attached recordings and examined the firing properties of ACh interneurons in acute striatal slices obtained from eIF4E Tg mice and wild-type littermates. ACh interneurons were identified by their distinctive large cell soma and their ability to generate spontaneous action potentials, which is maintained in striatal slices^67,68^. Our analysis revealed no differences in the firing frequency (Figure 4a and b) and coefficient of variation (Figure 4a and c) between the eIF4E Tg and their wild-type littermates, suggesting the absence of significant defects in the spontaneous activity of ACh interneurons.

**Figure 4.**
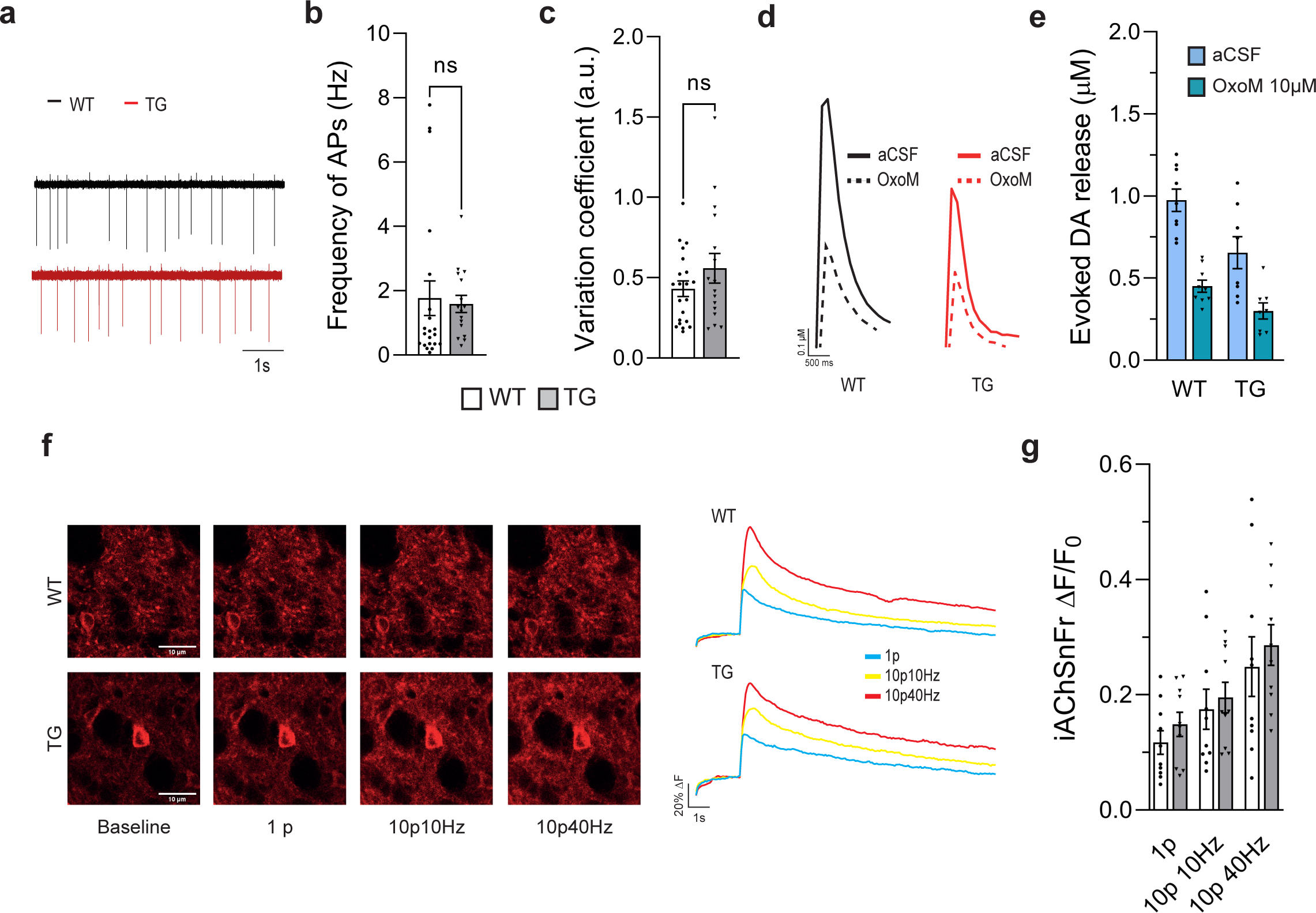
Preserved excitability, presynaptic response, and release in striatal ACh interneurons of the eIF4E Tg mice. (**a-c**) Cell-attached recording of ACh interneurons in acute corticostriatal slices. (**a**) Representative cell-attached traces of recordings presented in (**b** and **c**); wildtype (WT, black) and eIF4E Tg (TG, red). (**b**) Frequency of action potentials (AP) in Hz. Unpaired, two-tailed *t-*test, t_35_=0,3, ns. (**c**) Coefficient of firing variation (variation coefficient). Unpaired, two-tailed *t-*test, t_35_=1,3, ns. N= 20-17 cells/ 4-5 mice/ genotype. (**d-e**) FSCV recordings in the presence of oxotremorine M (OxoM 10 µM), an agonist of muscarinic receptors, in the dorsal striatum of acute corticostriatal slices. (**d**) Representative traces of FSCV recordings presented in (e); wildtype (WT, black) and eIF4E Tg (TG, red) before (aCSF, continuous lines) and after (OxoM, dotted lines) bath application of OxoM. (**e**) Peak concentrations of striatal evoked DA release recorded with FSCV in the presence of OxoM (dark blue) or vehicle (aCSF; light blue) following a single pulse stimulus. Two-way RM ANOVA, Genotype: F_(1, 15)_=8.4, p=0.01; Treatment: F_(1,15)_=92.5, p<0.0001; interaction Genotype:Treatment: F_(1, 15)_=3.4, p=0.09, ns. N= 10-12 slices/ 3 mice/ genotype. (**f-g**) Change of iAChSNFr fluorescence in response to ACh released by increasing electrical stimuli in acute corticostriatal slices. (**f**) Representative images and traces depicting iAChSNFr fluorescence at baseline and following 1p (blue), 10p10Hz (yellow) and 10p40Hz (red). (**g**) Variation of iAChSNFr fluorescence following 1 pulse (1p), 10 pulses at 10 Hz (10p10Hz) and 40 Hz (10p40Hz) detected with 2P microscope. Two-way RM ANOVA, Genotype: F_(1, 18)_=0.4, p=0.5, ns; Stimulation: F_(1, 19,34)_=54.4, p<0.0001; interaction Genotype:Stimulation: F_(2, 36)_=0.2, p=0.8 ns. N= 10 slices/ 2 mice/ genotype. For all graphs, bars represent group averages expressed as mean ± SEM and dots represent values for individual slices. ns= not significant.

ACh release is inhibited by presynaptic muscarinic receptors^69,70^. To determine if altered inhibitory feedback contributes to the diminished DA release observed in eIF4E Tg mice, we measured electrically evoked DA release with FSCV in the presence of the muscarinic receptor agonist oxotremorine M (OxoM). Application of OxoM (10 µM) to striatal slices caused a significant reduction of evoked DA release in wildtype and eIF4E Tg mice (Figure 4d and e; OxoM-sensitive DA release amounted to WT=0.52±0.06 µM and TG=0.35±0.06 µM; unpaired, two-tailed *t-*test, t_15_=1,84, n.s.), indicating that the function of presynaptic muscarinic receptors is intact in the eIF4E Tg mice.

Our results so far could be attributed to a deficit in stimulus evoked ACh release in eIF4E Tg mice. To explore this possibility, we first performed a colorimetric enzymatic assay to quantify ACh and choline concentrations in the striata dissected from the eIF4E Tg mice and wildtype controls^71^. Our results showed no significant differences in normalized ACh, and total choline concentration between the wild-type control and the eIF4E Tg mice (Figure S3a and b)

Next, we employed iAChSnFr, an optical sensor that binds to ACh and provides a means for measuring synaptic ACh release in acute striatal slices with two photon (2P) microscopy^72^. An AAV carrying the iAChSnFr was injected into the striatum of the eIF4E Tg mice and wild-type littermates. Striatal slices prepared at least 3 weeks post-surgery exhibited detectible iAChSNFr in the striatum, which responded to exogenously applied ACh with a dose-dependent increase in fluorescence (Figure S3c). Importantly, we observed no genotype-specific differences in the sensitivity of iACSnFr for bath applied ACh (50 µM) (Figure S3d and e).

We employed a series of electrical stimulations to induce ACh release in acute striatal slices and found no differences in the relative fluorescence of iACSnFr between eIF4E Tg mice and their wild-type littermates at any of the stimulation patterns tested (Figure 4f and g). These results indicate that ACh release elicited by electrical stimulation with similar frequencies to those used to elicit DA release in the FSCV experiments does not differ between eIF4E Tg and wildtype littermates.

Our results collectively demonstrate that the diminished DA release observed in the eIF4E Tg mice, while attributed to deficits in ACh signaling, is independent of ACh neurotransmission in the striatum.

### Impaired Ca2+ dynamics in the DA axons of the eIF4E Tg mice

When ACh binds to nAChR, it activates both sodium (Na+) and calcium (Ca^2+^) conductances^73^. This, in turn, initiates axonal action potentials followed by the opening of voltage gated Ca^2+^ channels and neurotransmitter release^61^. Given the direct association between axonal Ca^2+^ levels and neurotransmitter release, we employed an independent measure of presynaptic function by selectively expressing the calcium sensor GCaMP6s (GCaMP) in the nigrostriatal terminals. To this end, we injected an AAV carrying the *Cre*-dependent GCaMP gene into the substantia nigra of DAT-*Cre*/WT and DAT-*Cre*/TG littermates^68,74^. Three weeks later, we imaged fluorescent GCaMP transients in DA axons in acute striatal slices using 2P microscopy.

As GCaMP is not a ratiometric sensor for Ca^2+74^, we expressed the changes in fluorescence as a ratio between a train and a single pulse stimulation (ΔF/F_0_ train/ ΔF/F_0_ single pulse, referred to as ΔF/F_0_ ratio). Single pulse stimulation triggers a baseline GCaMP fluorescence response, while the train stimulation (10 pulses at 10 Hz; 10p10Hz) provides the GCaMP signal corresponding to a saturated DA release. As predicted, in slices from wild-type animals, we observed brighter Ca^2+^ transients in response to the train stimulation compared to a single pulse in aCSF, as shown by a ΔF/F_0_ ratio >1 between the change in fluorescence elicited by the train as compared to the single pulse (Figure 5a and b). A similar stimulation-dependent facilitation in Ca^2+^ influx was observed in slices from eIF4E Tg mice. Thus, like DA release measured with FSCV (see Figure 2a and c), the eIF4E Tg mice exhibit an intact frequency-dependent increase in presynaptic Ca^2+^ influx.

**Figure 5.**
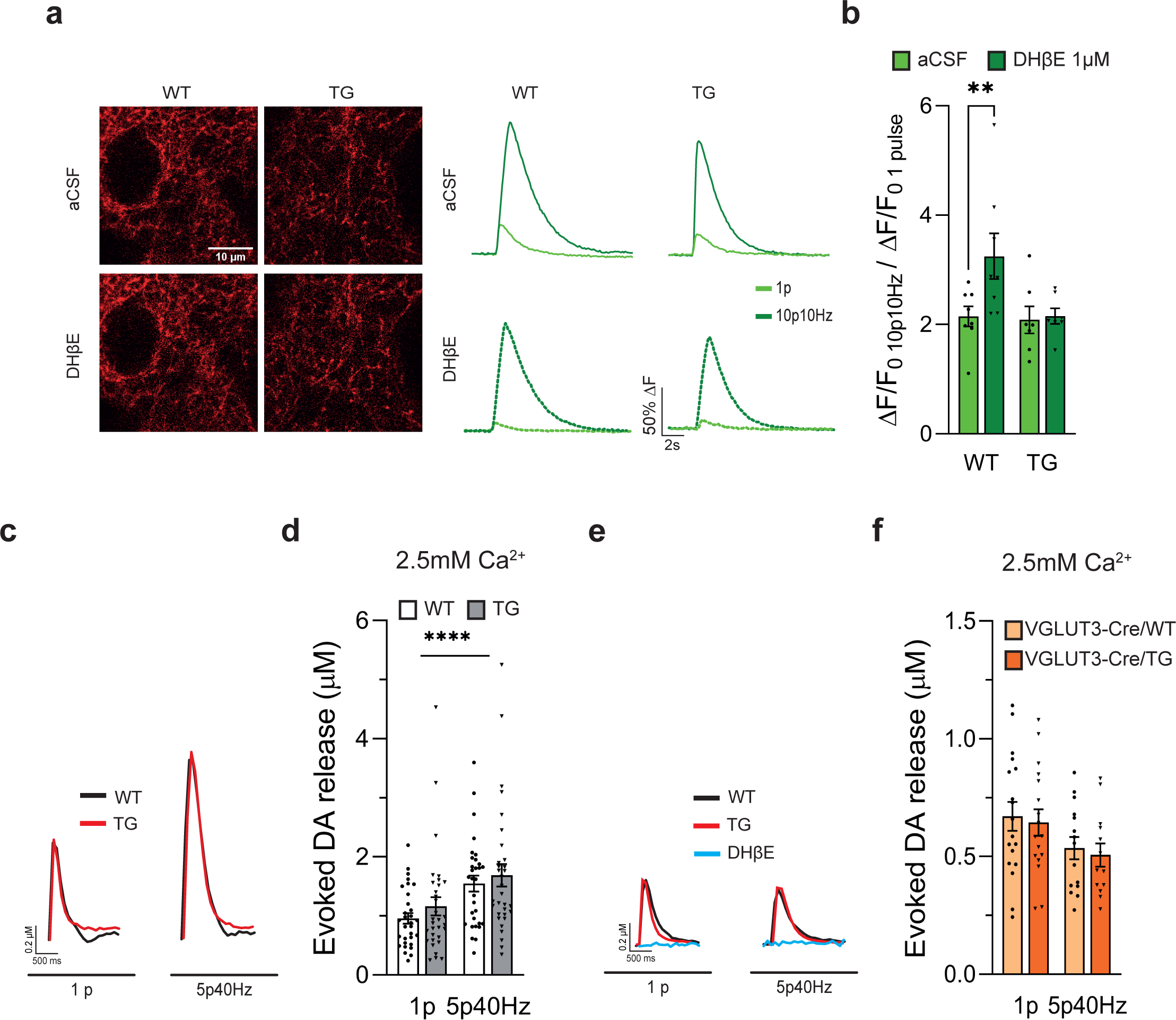
Altered activity dependent presynaptic Ca^2+^ dynamics triggered by ACh and normalized DA released in high extracellular Ca^2+^ in the eIF4E Tg mice. **(a-b)** Variation of GCaMP fluorescence in DA axons in response to different electrical stimuli in the presence of DHβE (10 µM) or vehicle (aCSF). (**a**) Representative images and traces of GCaMP fluorescence following 1p (light green) and 10p10Hz (dark green) in aCSF (solid lines) and DHβE (dotted lines) obtained with a 2P microscope. (**b**) Changes of GCaMP fluorescence following 10 pulses at 10 Hz (10p10Hz) versus 1 pulse (1p) in the presence of DHβE (dark green) or vehicle (aCSF; light green). RM Two-way RM ANOVA, Genotype: F_(1,13)_= 3.3, p=0.08, ns; main effect of Treatment: F_(1,13)=_6.7, p<0.05; interaction Genotype:Treatment: F_(1,13)_=4.8, p<0.05; **p<0.00, Multiple comparison’s post-hoc test. N= 7-8 slices/3 mice/ genotype. (**c-f**) FSCV recordings in the dorsal striatum of acute corticostriatal slices in higher extracellular Ca^2+^ (2,5 mM) aCSF. DA release was evoked by electrical (**c-d**) and optogenetic (**e-f**) activation of ACh interneurons (see Figure 3e-f). (**c**) Representative traces of FSCV recordings presented in (**d**); wildtype (WT, black) and eIF4E Tg (TG, red). (**d**) Peak concentrations of evoked DA release in 2,5 mM Ca^2+^ aCSF following a single pulse (1p) and 5 pulses at 40 Hz (5p40Hz) electric stimuli. Two-way RM ANOVA, Genotype: F_(1, 61)_=0.7, p=0.4, ns; Stimulation: F_(1, 61)_=152,9, p<0.0001; interaction Genotype:Stimulation: F_(1, 61)_=0.5, p=0.5, ns. N= 26-27 slices/ 6 mice/genotype. (**e**) Representative traces of FSCV recordings presented in (f); wildtype (WT, black) and eIF4E Tg (TG, red) and DHβE (blue). (**f**) Peak concentrations of evoked DA release induced by optogenetic activation of ACh interneurons (see Figure 3e-f) in 2,5 mM Ca^2+^ aCSF following a single pulse (1p) and 5 pulses at 40 Hz (5p40Hz) light stimuli. Two-way RM ANOVA, Genotype: F_(1,28)_=2.9, p=0.1, ns; main effect of Stimulation: F_(1,28)_=0.1, p=0.7, ns; Genotype:Stimulation: F_(1,28)_=0.3, p=0.5., ns. N= 17-13 slices/ 3 mice/ genotype. For all graphs, bars represent group averages expressed as mean ± SEM and dots represent values for individual slices. ns= not significant.

To investigate the role of β2-nAChR in axonal presynaptic Ca^2+^ transients, we then repeated this protocol in the presence of DHβE (1 µM). In acute striatal slices from wildtype mice, bath application of DHβE, which reduces DA efflux to a single pulse stimulation (see Figure 3a and b), resulted in a significantly higher train to single pulse (ΔF/F_0_) ratio compared to aCSF conditions (Figure 5a and b). By contrast, we did not observe any effect of DHβE in the eIF4E Tg mice (Figure 5a and b). Thus, our results show that eIF4E Tg mice exhibit a reduction in DA release that is concomitant with an impaired modulation of presynaptic Ca^2+^ influx by nAChRs.

To explore this further, we manipulated the driving force for Ca^2+^ by increasing its extracellular concentration from 2 to 2.5 mM. We then conducted FSCV in acute striatal slices of eIF4E Tg and wildtype mice and induced DA release with a single pulse and a train (5p40Hz). In elevated extracellular Ca^2+^, we did not observe genotype differences in response to any of the stimulations tested (Figure 5c and d), indicating that higher extracellular Ca^2+^ levels compensate for the altered β2-nAChR-mediated release of the eIF4E Tg mice, leading to a normalized DA release.

Finally, we investigated whether a similar increase in extracellular Ca^2+^ could normalize ACh-dependent DA release. We explored this possibility by measuring DA release with FSCV during optogenetic activation of ACh interneurons in striatal slices obtained from VGLUT3-*Cre*/WT and -/TG mice, as outlined in Figure 3e and f. In these experimental conditions, we observed an unaltered DA release in the eIF4E Tg mice (Figure 5e and f). Thus, higher levels of extracellular Ca^2+^ increase the release probability similarly in both genotypes, effectively bypassing the impaired β2-nAChR signaling in eIF4E Tg mice.

## Discussion

Our investigation revealed a reduction in DA release in the striatum of the eIF4E Tg mice, which can be ascribed to compromised functionality of β2-nAChR on DA axons. Specifically, optogenetic stimulation of striatal ACh neurons, in contrast to direct activation of DA terminals, unmasked an impaired evoked DA release in the eIF4E Tg mice. Importantly, this reduction occurred without concurrent impairments in morphology, biochemistry, and firing activity of DA neurons, as well as the firing activity and release patterns of striatal ACh interneurons assessed through a combination of electrophysiology recordings, 2P imaging and biochemical analysis. The reduced DA release was accompanied by behavioral inflexibility in the four-choice odor discrimination task, providing a link between ASD symptomatology and striatal DA dysfunction.

Correlative genetic and brain imaging studies in individuals with ASD provide support for the involvement of altered DA neuromodulation in the impaired executive functions associated with the disorder^6-8,30,31,34,35^. One such function altered in ASD is cognitive flexibility, defined as the ability to adapt one’s behavior according to a changing environment^30,31,33-35^. This dimension can be evaluated in ASD patients and preclinical models within a laboratory setting through reversal learning tasks^33^.

In our study, we employed the four-choice odor discrimination task^75,76^, an intradimensional reversal test sensitive to impairments in DA neuromodulation^36,37^. For instance, mice lacking the transforming growth factor β type II (TGF-βII), which display morphological alterations, E/I imbalance and altered firing pattern of DA neurons, also exhibit deficient reversal learning in the four-choice odor discrimination task^36^.

The eIF4E Tg mice exhibited impaired reversal learning but intact discrimination learning. Notably, during the reversal phase, the eIF4E mice made more perseverative errors, reflecting an inability to flexibly adapt and update a previously learned behavioral strategy. This aligns with our previous studies and those of others, demonstrating that eIF4E Tg mice and other preclinical ASD models characterized by hyperactive mTOR signaling exhibit impaired reversal learning but intact task acquisition accompanied by altered DA signaling^9,27,29,37,77^. For instance, a mouse model of Tuberous Sclerosis Complex 1 (TSC1), with selective genetic deletion of *tsc1* in DA neurons, showed impaired reversal learning in this task, along with reduced striatal DA release detected by FSCV in acute corticostriatal slices^37^, similar to the eIF4E Tg mice. However, unlike the eIF4E Tg mice, DA neurons lacking *tsc1* displayed extensive morphological and physiological impairments^37^.

TSC1, which forms a complex with TSC2, serves as a negative regulator of mTOR signaling^78,79^. Consequently, the genetic deletion of *tsc1* results in constitutive hyperactive mTOR signaling, leading to augmented neuronal protein synthesis (via eIF4E) and, concurrently, reduced macroautophagy^14,80,81^. Although the specific molecular details are yet to be elucidated, it is plausible that the genetic deletion of *tsc1* in DA neurons disrupts axonal release, morphology, and cell body physiology due to changes in both mTOR-dependent translational and macroautophagy. This hypothesis gains further support from the abnormal morphology observed in DA neurons of mice with a conditional genetic deletion of *atg7*, a critical regulator of macroautophagy^82^. The morphological alterations in these neurons characterized by an increased dimension of the soma and axon terminals^82^, closely resemble those observed in DA neurons lacking *tsc1*. Conversely, the overexpression of *eIF4E*, and consequently, the increased cap-dependent translation^27^, results in more nuanced alterations in DA neuron physiology, particularly in the regulation of DA release. Although not accompanied by morphological alterations, these changes are sufficient to induce behavioral inflexibility.

Alterations in DA neuromodulation and behavioral inflexibility have also been reported in a mouse model of fragile X syndrome (FXS). FXS is a genetic form of intellectual disability and ASD characterized by the genetic inactivation of *Fmrp*, encoding for an RNA-binding protein that represses the translation of a specific subset of mRNAs associated with eIF4E. Thus, FXS model mice exhibit increased eIF4E-dependent protein synthesis along with hyperactive mTOR signaling^83,84^. The dysregulated protein synthesis is accompanied by diminished evoked DA release detected by FSCV in acute corticostriatal slices^85^ and behavioral inflexibility in a touchscreen operant task^86^. Although the synaptic mechanisms leading to impaired striatal DA release in FXS remain unclear, it is tempting to speculate that the synaptic and behavioral similarities between eIF4E Tg and FXS mice result from increased translation of overlapping mRNAs. Indeed, a fraction of the eIF4E-sensitive transcripts, preferentially translated in eIF4E Tg mice may be repressed by FMRP, and thus specifically translated in FXS.

Our data also demonstrate that the DA release deficits in the eIF4E Tg mice results from a malfunction of axonal β2-nAChR. Importantly, a4β2-subunit containing nAChR, which are expressed in DA terminals, are reduced in postmortem brain tissues of ASD patients, with no corresponding changes in mRNA levels, suggesting a post-transcriptional modification of these ion channel subunits^87^. Mice lacking the β2 subunit of nAChR display deficits in executive functions reminiscent of the perseverative behaviors observed in the eIF4E Tg mice and ASD patients^88,89^. Interestingly, these mice display reduced DA release at various stimulation frequencies while maintaining activity-dependent release dynamics^90^. These previous findings align with our own results, indicating that an altered β2-nAChR function would lead to a sustained reduction in DA release while maintaining activity-dependent release dynamics, similar to what is observed when eIF4E is overexpressed.

We further explored the role of β2-nAChR in activity-dependent release dynamics by employing an independent measure of presynaptic function based on the strong functional coupling between presynaptic Ca^2+^ levels and neurotransmitter release. To this end, we selectively expressed GCaMP6 in DA axon terminals and measured presynaptic Ca^2+^ transients in response to variation of stimulation frequency^74^. β2-nAChR inhibition has been demonstrated to reduce the release probability at DA synapses, thereby facilitating release during high frequency stimulations^59,60,91,92^. Consistently, presynaptic Ca^2+^ transients show a similar facilitation in the presence of DHβE in wildtype mice (see Figure 5a and b), suggesting that β2-nAChR blockade reduces the release probability at DA synapses by altering presynaptic Ca^2+^ dynamics. In contrast, the DHβE-induced facilitation of presynaptic Ca^2+^ transients was reduced in slices from the eIF4E Tg mice (Figure 5a and b). This indicates that the DA synapses of the eIF4E Tg mice are less sensitive to β2-nAChR inhibition, exhibiting impaired facilitation of presynaptic Ca^2+^ transients, thereby maintaining a high release probability. In summary, these findings indicate that the high-pass filter function provided by axonal β2-nAChR on DA synaptic transmission is compromised in the eIF4E Tg mice.

Recent findings suggest that DHβE abolishes action potentials in DA axons induced by optogenetic activation of ACh interneurons, implying a direct role for β2-nAChR in both axonal depolarization^61^ and release^59,60,90-92^. These two effects could be interconnected yet independent, with depolarization depending on the influx of both Na^+^ and Ca^2+^, while the release relying solely on Ca^2+^. Our imaging and electrochemical results do not allow us to distinguish between these two effects of β2-nAChR function and further studies are required to determine whether only release, or both axonal action potentials and release, are impaired in the eIF4E Tg mice.

While our data points to an altered function of β2-nAChR, which are expressed in DA axons, the underlying molecular mechanisms remain speculative. It is known that agonist-binding rapidly desensitizes nAChR, with kinetics significantly influenced by the subunit composition ^93,94^. In neurons, eIF4E regulates the translation of specific “eIF4E sensitive” mRNAs that harbor specific secondary structure elements in the 5’ UTR^14,95-98^. Therefore, it’s conceivable that β2-subunits may be preferentially translated when eIF4E is overexpressed, resulting in nAChR with altered subunit compositions and impaired desensitization dynamics. In line with this speculation, recent findings have demonstrated a selectively high expression of CHRNB2, which encodes the β2 subunits of nAChR, in patient-derived induced pluripotent stem cells (iPSCs) from individuals with FXS^99^.

Alternatively, nAChR with altered subunit compositions may not be trafficked to the membranes or could be misplaced, thereby leaving DA axons with aberrant nAChR. Moreover, mRNA encoding for other nAChR subunits expressed in DA neurons, such as α4 and 6^92,100,101^, may also be sensitive to eIF4E overexpression, contributing to impaired nAChR. Future studies are necessary to elucidate how eIF4E overexpression alters the translatome of DA neurons, including translation of nAChR subunits leading to altered assembly and trafficking.

The absence of a high-pass filter provided by β2-nAChR, as revealed by our study, may offer a mechanistic explanation for understanding behavioral inflexibility in ASD. Indeed, β2-nAChR-triggered DA release is suggested to play a crucial role in amplifying physiologically relevant stimuli encoded by the various firing modes of DA neurons (i.e., tonic vs burst firing)^45,47,52,102^. An aberrant β2-nAChR filtering system as in the eIF4E Tg mice, would not only hinder the dynamic probability of DA release in response to neuronal activity. It may also lead to a desynchronization of ACh interneurons since these interneurons respond directly to altered striatal DA levels via D2 receptors^102,103^.

Behaviorally, the absence of dynamic amplification in DA levels and ACh desynchronization in response to salient stimuli may lead to an aberrant reward-prediction error, causing the eIF4E Tg mice to perseverate with an acquired behavioral strategy even if it is no longer reinforced.

Given the crucial significance of the coincident activity of DA and ACh neurotransmission in the striatum for the accurate computation of a reward-prediction error, pharmacological approaches aimed at indiscriminately elevating striatal DA levels might not adequately improve the cognitive aspects of repetitive and perseverative behaviors in ASD.

In conclusion, our study demonstrates that in the eIF4E overexpression model of ASD, striatal DA neurotransmission is hindered by aberrant functionality of β2-nAChR on DA axons. These findings provide valuable insight into the intricate interplay between ACh and DA in striatal-dependent functions, highlighting the potential cellular origin of behavioral inflexibility present in ASD.

## Acknowledgments

We extend our gratitude to Gilad Silberberg, Abdel El Manira, Gilberto Fisone, Konstantinos Meletis, Alessandro Usiello, Sten Grillner, and all the members of the Fisone, Borgkvist and Santini labs for their invaluable methodological assistance and insightful discussions. Special thanks to Patrik Ernfors and Gilberto Fisone for generously providing us with the founders of vGLUT3- and DAT-*Cre* colonies, respectively. We also thank Rodrigo Espana for sharing the single card version of the Demon Voltammetry Software, Qian Yu and the KI Animal Behavioral Core Facility for the support received in conducting the behavioral study, Kristoffer Tenebro Berglund, the staff, and the veterinarians of the Comparative Medicine Biomedicum (KM-B) for their continuous support and assistance with the maintenance of the mouse colonies. This work was supported by the Knut and Alice Wallenberg Foundation (Wallenberg Academy Fellow Grant KAW 2017-0169 and project grant 2020-0054 to E.S.), the Swedish Research Council (2016-02758 to E.S. and 2016-03129 to A.B.), the Stiftelsen Olle Engkvist Byggmästare (E.S. and A.B.), Åhlén’s foundation (A.B.), Magn. Bergvall’s foundation (A.B.), The Strategic Research Program in Neuroscience (StratNeuro) starting (E.S. and A.B.) and bridging (E.S.) grants, Karolinska Institute starting (E.S.) and KID (E.S. and J.C.R.) grants. D.S. and E.M. were supported by NIDA R01DA07418 and JPB Foundation. O.J.L. was supported by NIH F30 MH114390.

## Author contributions

Conceptualization, O.J.L., A.B. and E.S.; Methodology, A.B. and E.S.; Validation, A.B. and E.S.; Formal Analysis, J.C.R., A.A., A.B. and E.S.; Investigation, J.C.R., A.A., A.M., E.M., J.K., O.J.L., A.B. and E.S.; Resources, A.B. and E.S.; Writing - Original Draft, A.B. and E.S.; Writing - Review & Editing, J.C.R., A.A., E.M., J.K., O.J.L., D.S., A.B. and E.S.; Visualization, J.C.R., A.B. and E.S.; Supervision, A.B. and E.S.; Project Administration, A.B. and E.S.; Funding Acquisition, A.B. and E.S.

## STAR Methods

### Animals

eIF4E*^wt/_β_tEif4e^* ^27,104^, VGLUT3-*Cre^wt/+^* (Tg(Slc17a8-icre)1Edw/SealJ; JAX Strain#018147) and DAT-*Cre^wt/+^* (Slc6a-Cre knockin)^62,63^ mice (referred to as eIF4E Tg, VGLUT3-*Cre* and DAT-*Cre*, respectively) were previously generated as described in^27,104,62,63,27,66,104,^.

Mice were sex-separated and group-housed (up to 5 mice for cage) in a temperature (23C) and humidity (55%) controlled environment, on a 12-h light/dark cycle with water and food available *ad libitum*. Only male mice not older than 4 months were used for the experiments ^27^.

All animal experiments were compliant with the ethical permit issued by the Swedish Board of Agriculture (Ethical number: 18194-2018) and were performed in accordance with the European Parliament and Council Directive 210&63/EU, 22nd September 2010 for experimentation animals’ protection.

### Breeding strategy

The DAT-Cre, VGlut3-Cre and eIF4E Tg mice were maintained in hemizygosis by crossing them with C57BL/6J mice purchased by Janvier. Double transgenic mice were generated by crossing DAT-*Cre^wt/+^* or VGLUT3-*Cre^wt/+^* with eIF4E*^wt/_β_tEif4e^*. The offspring expressing *Cre* in DA neurons or striatal acetylcholine interneurons, respectively along with either overexpressing (DAT-*Cre^wt/+^*/ eIF4E*^wt/_β_tEif4e^* or VGlut3-*Cre^wt/+^*/ eIF4E*^wt/_β_tEif4e^* referred to as DAT-Cre/eIF4E and VGlut3-Cre/eIF4E, respectively) or not (DAT-*Cre^wt/+^*/ eIF4E*^wt/wt^* or VGlut3-*Cre^wt/+^*/ eIF4E*^wt/wt^* and referred to as DAT-Cre/WT and VGLUT3-Cre/WT) eIF4E were utilized in this study. All mice were maintained in a C57BL/6J background.

### Genotyping

Genotyping was done by PCR; genomic DNA was obtained from earmarking biopsies. Tissue samples were digested overnight (minimum 6h) at 56°C, shaking at 1000-1100 rpm, in 400μL of the following buffer: 100mM Tris-HCL pH 7.5, 1mM EDTA, 250mM NaCL, 0.2% SDS, Proteinase K (Invitrogen) 0.1mg/mL. Afterwards, lysates were diluted 1:10 in dH2O and used as template for PCR reaction carried out using the KAPA2G mastermix (Kapa Biosystems). Primers used in eIF4E transgene amplification were: 5’-CACAGCTACAAAGAGCGGCTCCACC -3’ and 5’-CACTGCATTCTAGTTGTGGTTTGTCC-3’. Those used for DAT-Cre amplification were: 5’-CATGGAATTTCAGGTGCTTGG-3’, 5’-ATGAGGGTGGAGTTGGTCAG-3’,5’-CGCGAACATCTTCAGGTTCT-3’, and for vGLUT3-Cre amplification: 5’-ACA CCG GCC TTA TTC CAA G-3’, 5’-AGA TGT CTT ATG GAG CCA CCA C-3’, and 5’-CTG AGA CCA AGG TCC ATA TTC C-3’.

### Acute Brain Slices

For FSCV experiments, brain slices were prepared as previously shown ^105^. Briefly, mice underwent cervical dislocation, and the brain was removed and placed on a high sucrose cutting solution, previously cooled to 4°C: 10mM NaCl, 2.5mM KCl, 25mM NaHCO_3_, 0.5mM CaCl_2_, 7mM MgCl_2_, 1.25mM NaH_2_PO_4_, 180mM sucrose, 10mM glucose; bubbled with 95% O_2_/5% CO_2_ to pH 7’4. Brains were mounted on a VT1200 vibratome (Leica Biosystems) and coronal sections (250 μm) including the striatum were collected. Slices were transferred to Artificial Cerebro-spinal Fluid (aCSF) containing (in mM): 125 NaCl, 2.5 KCl, 25 NaHCO_3_, 2 CaCl_2_, 1 MgCl_2_, 1.25 NaH_2_PO_4_, and 10 glucose bubbled with 95% O_2_/5% CO_2_ to pH 7.4, at 34°C for 30 minutes. After that, the slices were kept in oxygenated (95% O_2_/5% CO_2_) aCSF at room temperature.

For Patch-Clamp electrophysiology, and 2P Microscopy acute brain slices were prepared and maintained using the two-step *N-*methyl-D-glucamine (NMDG) based protective recovery method^106^. Slices were initially prepared using NMDG-based solution containing (in mM): 92 NMDG, 30 NaHCO_3_, 2.5 KCl, 20 HEPES, 2 thiourea, 1.25 NaH_2_PO_4_, 3 Na-pyruvate, 5 ascorbic acid, 25 glucose, 0.5 CaCl_2_ and 10 MgCl_2_ titrated to pH 7.4. During the experiment, the slices were stored in the HEPES-based holding solution containing (in mM): 92 NaCl, 30 NaHCO_3_, 2.5 KCl, 20 HEPES, 2 thiourea, 1.25 NaH_2_PO_4_, 3 Na-pyruvate, 5 ascorbic acid, 25 glucose, 2.5 CaCl_2_ and 2 MgCl_2_ titrated to pH7.4. Mice underwent cardiac perfusion with 25ml chilled NMDG solution prior to brain removal and dissection. Brains were mounted on a VT1200 vibratome (Leica Biosystems) and coronal sections (250 μm) containing the striatum or the substantia nigra were collected and stored in NMDG-based solution for 10 minutes at 32°C. Subsequently, the slices were maintained in HEPES-based holding solution at room temperature. All solutions were oxygenated with 95% O_2_ and 5% CO_2_.

### Fast-Scan Cyclic Voltammetry (FSCV)

Carbon fibre microelectrodes (CFMs) were made in house as follows: 7μm thick carbon fibres (Good Fellow, Huntingdon, GB) were inserted into 0.69x1.20x100mm borosilicate glass tubes (Science Products). Tubes were pulled to seal the glass around the fibre, and the exposed fibre was cut ∼150 μm length. To improve the seal around the carbon fibre, CFMs tips were immersed in melted paraffin, and extra coating was removed with xylene^107^. CFMs were calibrated after each recording by quantifying DA at known concentrations (1.25, 2.5, 5 μM) as previously described ^108^.

Striatal slices were placed in the submersion recording chamber of the rig and continuously perfused with oxygenated (95% O_2_/ 5% CO_2_) aCSF at 32-34°C for at least 10 min before starting the recordings. CFMs filled with 1M KCl, were inserted in the dorsolateral striatum and received a triangular voltage wave (-0.4V to +1.2V at 400V/s) every 100 ms. The resulting currents were measured with a Chem-clamp 5MEG amplifier (Dagan corporation, Minnesota, USA). Slices were stimulated with a bipolar stainless-steel electrode placed ∼150μm from the recording electrode. Using an Iso-Flex stimulus isolator (A.M.P.I.), single pulses (1ms × 300μA) were applied every 90s to determine a stable DA release. For experiments involving train stimulations, 250s were allowed between stimuli. Recordings and data quantification were done with the Demon Voltammetry suite software ^39^.

All drug inhibitors were dissolved in water, unless otherwise stated and bath applied for at least 15 minutes after achieving a stable baseline. We employed the following drugs: 1μM DHBE (Dihydro-β-erythroidine hydrobromide, Tocris), 10 μM Oxotremorine (Tocris), 2μM Sulpiride (Sigma; dissolved in DMSO). The concentrations employed in these studies are in a range compatible with previously published FSCV experiments ^58,59,69^.

When modified aCSF was used, cations concentrations were adjusted to maintain osmolarity: from 2mM Ca^2+^ /1mM Mg^2+,^ to 2.5mM Ca^2+/^0.5mM Mg^2+.^

In FSCV recordings with optogenetic stimulation, optical probe was placed ∼150μm from the recording electrode, and 760nm light was delivered in 3ms pulses every 120s.

### Patch-Clamp Electrophysiology

Recordings were obtained using a patch clamp electrophysiology rig fitted with SciCam pro camera (Scientifica, United Kingdom) equipped with a 40 × 0.8 NA water-immersion objective (LUMPlanFLN, Olympus, United States) and Dodt contrast tube optics. Recordings were obtained with a Multiclamp 700B amplifier (Molecular Devices) and Axon Digidata 1550B digitizer (Molecular Devices, United States), using pCLAMP 11 software (Molecular Devices, United States). Data were low-pass filtered at 10 kHz. For all recordings, borosilicate glass capillaries were used to prepare electrodes with 3-4MOhm tip resistance. Slices were maintained in a submersion recording chamber with oxygenated (95% O2/ 5% CO2) aCSF supplied at 3ml/min and maintained at 32-34°C.

Extracellular recordings of cholinergic interneurons

Recording electrodes were filled with aCSF to avoid liquid-junction potential. Putative cholinergic interneurons were identified based on their morphology and chosen at a depth of at least 50µM into the slice, due to their relatively big soma size. During recording, a weak seal (< 10 times pipette resistance) was maintained as to avoid membrane perturbations that may influence tonic firing activity. Spontaneous firing activity of the cells was recorded for at least 10 minutes, and seal resistance was monitored during the whole procedure.

### Intracellular and extracellular recordings, identification of DA neurons

Recording electrodes were filled with potassium-gluconate based internal solution containing (in mM): 120 K-gluconate, 20 KCl, 4MgATP, 0.3Na2GTP, 5 Na2-phosphocreatine, 0.1 EGTA and 10 HEPES at pH 7.25 and osmolarity ∼300mOsm. Neurobiotin (Vector Labs, 0.2%) was added to the internal solution for post-hoc identification.

Dopaminergic cells were initially identified based on size as well as firing properties upon entering whole-cell configuration. Prior to breaking through the membrane, cells were held in cell-attached configuration with voltage held at -70mV for one minute to measure extracellular firing. The rest of the recordings taken upon entering whole-cell configuration were performed in current-clamp mode. Ramp current injection was performed ranging from -300 to +700pA over 1 second with 1 second intersweep interval. Following that we performed stepwise current injection from -300pA to +700pA in 50pA increments. Each current step was held for 500ms with 500ms intersweep interval. Access resistance (Ra) was measured before and after the current injection recordings in voltage-clamp mode and recordings were discarded if Ra exceeded 30MOhm or changed by over 10%.

After recording, the slices were fixed overnight in 4% paraformaldehyde (PFA) in 0.1M PBS (pH 7.4) at 4°C and underwent immunofluorescence staining to identify biotin-filled DA neurons ^105^. Briefly, following a series of washes in tris-buffered saline (TBS), slices were incubated in Streptavidin Alexa Fluor™-647 (Invitrogen, 1:200) in 0.6% Triton X-100 in TBS for 48 hours. Slices were then incubated for 1 hour in blocking solution containing 10% normal goat serum (NGS) in 0.6% TBS-TritonX100 followed by immunostaining for tyrosine hydroxylase (TH) which was used as a cellular marker for dopaminergic cells. Cells were incubated in 2% NGS / 0.6% TBS-Triton containing mouse anti-TH (Millipore, #MAB318, 1:1000) for 72 hours, followed by overnight incubation in 2% NGS / 0.6% TBS-triton with goat anti-mouse Alexa Fluor™ 488 (Invitrogen, #A32723, 1:500). Only recordings obtained from neurons positive for TH, and thus considered DA neurons, were included in this study.

Analysis of patch-clamp recordings

Analysis of patch-clamp data was performed using either ClampFit (Molecular Devices) or the open-access Python-based software Stimfit 0.15^109^. The extracellular firing frequency of DA and ACh neurons was determined by counting spikes in a 10-second or 60-second window, respectively, to determine the rate per second. Variation coefficient was calculated by dividing the standard deviation by the mean (VC= σ/µ), for each experimental group.

The current at which the first action potential fired with ramp current injection was used to determine the rheobase current. Action potential duration was determined using the half-width of the action potential, which measures the difference in time between the half-max amplitude during depolarisation and repolarisation. Current-frequency (IF) plots were generated from the action potential frequency (Hz) at each current step. Sag ratio was determined using the ratio between the steady state decrease in voltage and the peak decrease in voltage with the hyperpolarising current injection of -300pA. The voltage response to-100pA current injection was used to determine the input resistance (Ri) and an exponential line fit from the initial voltage change was used to determine the membrane time constant (⍰). Membrane capacitance was calculated using ⍰/Ri.

### 2 photon microscopy

Acute striatal slices were transferred to a recording chamber and allowed to stabilize for at least 15 min. After, a tungsten concentric bipolar electrode with a 3µm tip size (World Precision Instruments) was positioned on the surface of the slice, and images were captured approximately ∼150µm away from the stimulating electrode. Using two IsoFlex stimulus isolators (A.M.P.I.) connected in parallel, slices were stimulated with the following protocol (±250µA x 500µs): a single pulse, followed by 10 pulses at 10Hz, and another set of 10 pulses at 40Hz, with a 250s interval between stimulations. Fluorescent biosensors were excited with a Ti:sapphire laser (Ultra II, Coherent) and imaging was performed using a commercially available 2-photon microscopy setup (Scientifica) equipped with a 16x objective, 0.8NA (Nikon).

Axonal-GCaMP6s and iAChSnFr were excited at 920nm and 950nm respectively. Images were acquired with the following settings: 5x zoom, 512x512 px frame size, 1 frame/image (scan rate 1Hz).

Image quantification was performed using ImageJ, where fluorescence over time was analyzed. The baseline for fluorescence values was established by averaging frames before stimulation. Subsequently, maximal fluorescence values were normalized through baseline subtraction.

### Western Blot

Striatal sample preparation was performed as previously described^110^. Briefly, mice were rapidly decapitated, and their head was briefly placed in liquid nitrogen. Brains were subsequently removed and the striata were rapidly dissected and flash frozen in liquid nitrogen. The tissues were sonicated in 750μL of 1% SDS and the homogenates were boiled at 100°C for 10 min. The protein content of the homogenates was determined with the Pierce BCA Protein Assay kit. The samples were prepared by adding Lemli buffer (4X) and by boiling the samples for another minute. Equivalent amount of protein per sample (25-30 μg/well) were loaded into 10% polyacrylamide gels as previously described^110^. Proteins were transferred to Immobilon FL PVDF membranes (pore size 0.2 μm). Blots were blocked with a blocking solution composed of Odyssey Blocking Buffer:TBS + 0.1% Tween-20 (TBST) in a ratio 1:1 for one hour at room temperature. Blots were then incubated with primary antibody (Cell Signalling: DARPP-32, #: 2306S and actin, # 4970; Millipore: TH, #MAB318 and DAT, #MAB369; Sigma: vMAT2, #V9014); all diluted 1:1000) diluted in Odyssey Blocking Buffer overnight at 4 °C. Blots were then washed with TBST and incubated with secondary antibodies (IRDye, LICOR: anti-rabbit for DARPP-32, actin and vMAT2; and anti-mouse for TH; diluted 1:10000;) for one hour at room temperature. Blots were developed using the Odyssey imaging system (LICOR). Western blots were analysed in Image Studio Lite (LICOR).

### Ultra-high performance liquid chromatography tandem mass spectrometry **(UHPLC-MS/MS)**

Concentrations of DA, DOPAC and HVA in the striatal samples were measured by UHPLC-MS/MS following derivatization with benzoyl chloride as previously described^111,112^. Briefly, 20 µl volumes of the deproteinated supernatants from the tissue homogenates (1:10 w/v in MeOH) were mixed with 20 µl of the internal standard mixture (1 µM of each deuterated analyte standard DA-d4, DOPAC-d5, HVA-d3 in MeOH) followed with pipetting 20 µl benzoyl chloride (2% in ACN) and 20 µl sodium carbonate (100 mM in water, pH 10.5). The mixtures were vigorously shaken after each pipetting step. The resulting derivatized solutions were mixed for 5 min. The reaction was terminated by pipetting 20 µl sulfuric acid (1% in water), 10 µl of the final solution was injected on column. The UHPLC-MS/MS system included a Waters Xevo TQ-S micro triple quadrupole mass spectrometer with the electrospray ionization source operating in a positive mode, and an ACQUITY UPLC system (all purchased from Waters Corporation, Milford, MA, USA). The calibration curves were constructed in the range of 0.2 – 1000 ng/ml.

### Detection of striatal ACh content

ACh levels in the striatum were measured with Choline/Acetylcholine Assay kit (Abcam, #ab65345) following the colorimetric procedure ^71^. Briefly, mice were rapidly decapitated, and their head was briefly placed in liquid nitrogen. Brains were removed and the striata were rapidly dissected and flash frozen in liquid nitrogen. Each striatum was homogenized in 50 uL of ice-cold Choline Assay Buffer with a Dounce homogenizer. The homogenates were transferred to new tubes, centrifugated at 14,000 rpm for 10 min at 4C. An aliquot (10uL) of each sample was transferred to a new tube and utilized to measure the protein content with Pierce BCA Protein Assay kit. The remaining samples were subjected to the Choline/Acetylcholine assay as per manufacture’s instruction.

### Immunohistochemistry

Animals were deeply anesthetized with Pentobarbital, and transcardially perfused with 0.9% saline solution for one min followed by 4% paraformaldehyde (PFA) in phosphate buffer for 10 min. Brains were extracted and post-fixed overnight (same paraformaldehyde solution), cut in 30μm thick sections using a VT1200 vibratome (Leica Biosystems) and stored in cryoprotectant solution (30% Glycerol, 30% Ethylene glycol in 0.1M NaHPO_4_) at -20°C. Brain sections were processed for immunohistochemistry as follows: sections were washed three times in PBS for 10 minutes, permeabilized in 0.1% Triton-X100 in PBS (PBS-T), three times for 10 minutes, and incubated for 1 hour in blocking solution containing: 2% Normal Goat Serum (NGS), and 0.1% PBS-T. After blocking, sections were incubated overnight at 4°C in 2% NGS + 0.1% PBS-T containing a combination of the following antibodies: guinea pig anti-ChAT (Synaptic Systems, #297015, 1:500), mouse anti-Tyrosine Hidroxylase (Millipore, #MAB318, 1:100), chicken anti-GFP (Abcam, #Ab13970, 1:2000). After washing, sections were incubated for 2h with 2%NGS and 0.1%PBS-T containing a combination of the following secondary antibodies (Invitrogen, for all 1:500): goat anti-rabbit conjugated with Alexa 488 (#A11034), goat anti-chicken IgG conjugated with Alexa 488 (#A11039), goat anti-guineapig conjugated with Alexa 568 (#A11075)1:500). Sections were mounted, after washing, using a medium for fluorescent sections: Fluromount-G with DAPI (Invitrogen).

For Immunofluorescence quantification, sections were imaged at 20x with confocal microscope (Zeiss LSM800). To avoid biases in quantification, z-stacks at 1μm intervals across the vertical axis of the section were acquired. Afterwards, images were processed with ImageJ to compile the z-stacks into single images, and immunofluorescent cells were manually counted.

### Stereotactic surgeries

Mice were injected with buprenorfine (intraperitoneally (i.p.); 5mg/kg), anaesthesized with isoflurane (2%) and placed in a stereotaxic frame (Stoelting). Adenovirus (AAV) injection was performed using a glass pipette (Drummond, 15-20µm tip size). A fitted plunger controlled by a hydraulic injector (Quintessential Stereotaxic Injector, Stoelting), was inserted into the pipette, and used to inject the viral solutions (250nL and 100nL in striatum and SNc, respectively) at the steady rate of 50nL/min. We utilized the following coordinates (from bregma): Striatum (0.8 AP, 1.8ML, 2.4DV), SNc: (-3.1 AP, ML 1.25, DV 3). Rimadyl and buprenorphine (i.p., 0.1mg/kg) were administered twice a day for 3 days after surgery. The mice were left to recovery after surgeries for at least three weeks before being employed in other type of experiments.

AAVs injected in the striatum: pAAV-Ef1a-DIO EYFP was a gift from Karl Deisseroth (Addgene viral prep #27056-AAV5; http://n2t.net/addgene:27056; RRID:Addgene_27056), pAAV.hSynap.iAChSnFR was a gift from Loren Looger (Addgene viral prep # 137950-AAV1; http://n2t.net/addgene:137950; RRID:Addgene_137950)^72^, pAAV-EF1a-DIO-hChR2(H134R)-mCherry-WPRE-HGHpA was a gift from Karl Deisseroth (Addgene viral prep # 20297-AAV5; http://n2t.net/addgene:20297; RRID:Addgene_20297). Those in the SNc: pAAV-hSynapsin1-FLEx-axon-GCaMP6s was a gift from Lin Tian (Addgene viral prep # 112010-AAV5; http://n2t.net/addgene:112010; RRID: Addgene_112010)^113^, pAAV-EF1a-DIO-hChR2(H134R)-mCherry-WPRE-HGHpA was a gift from Karl Deisseroth (Addgene viral prep # 20297-AAV5; http://n2t.net/addgene:20297; RRID:Addgene_20297).

### Four-choice odor discrimination and reversal-learning paradigm

Adult male mice were subjected to the four-choice odor discrimination and reversal-learning paradigm ^36,37,75,76^ to assess learning and cognitive flexibility. The mice were food restricted with *ad libitum* access to drinking water and maintained at around 80% of their body weight until completion of the behavioral tests. The food restriction began 3 days before pre-training. The animal weights were recorded in the morning and food was given at the end of the day.

We found no significant differences in the body weight of wildtype and eIF4E Tg mice prior the test (WT: 30.8 ± 1.1 gr and TG: 29.4± 0.9 gr), during the three days of food restriction (e.g., weight at the last day of food restriction WT: 27.2 ± 0.9 gr and TG: 25.9 ± 0.9 gr) and the three days of the test (e.g., weight at the last day of the test WT: 25.2 ± 0,8 gr and TG: 24.1 ± 0.6 gr) as indicated by Two-way RM ANOVA, showing a main significant effect of the Time (F_(6, 72)_=91.5, p<0.0001) but not main significant effect of the genotype (F_(1, 12)_=1.1, p=0.3) and Time:Genotype interaction (F_(6, 72)_=0.2, p=0.9).

The test arena consists in an opaque white custom-made box (45 x 45 x 45 cm). Odour stimuli were presented in dark ceramic pots (measuring 7cm in diameter and 5cm in depth) located in each corner of the box. Pots were sham baited with Cheerios Honey cereals (Nestle’ Sverige AB, Helsingborg, Sweden) placed underneath wood shavings. The apparatus and the pots were cleaned with 70% ethanol and carefully washed with soap at the end of each testing day.

The first day of pre-training (day1; see schematic Figure 1a), mice were allowed to freely explore the testing arena and the pots. Around 1/8^th^ of Cheerios Honey was placed inside each empty pots, located in the four corners of the box. The mouse was placed in a central glass cylinder that was lifted to allow exploration and consumption of the cereals in the pots. After 10 min, the mouse was returned in the cylinder and the pots re-baited. This was repeated 3 times for a total habituation time of 30 minutes.

The second pre-training day was the shaping phase of the test (day 2) used to teach the mice to dig the food rewards (1/8^th^ of Cheerios Honey) buried in wood shavings. In the shaping phase, only one pot was employed with increasing amount of shaving material to gradually cover the food reward. The corner containing the pot was alternate in each trail and all corners were equally rewarded. Trials were untimed and consisted of 2 trials without wood shavings, two trials with a dusting of shaving, two trails with a quarter full pot, two trails with half full pot and four trials with the food reward completely buried.

On the odour discrimination and reversal tests (day 3) the wood shavings were freshly scented with organic essential oils (Örtagubben AB, Stockholm, Sweden). All the essential oils (100% pure) were diluted 1:10 in odourless mineral oil and mixed at 0.05 ml/gr of shavings. In both tests, the stimulus presentation was pseudo-randomized such that an odour was never in the same quadrant two trials in a row. In the discrimination and reversal tests *criterion* was met when the animal completed 8 out of 10 consecutive trials correctly.

During the discrimination phase the mice had to discriminate between four different odours (O1-4: thyme, clove, rosemary, and cardamom) and learn the one (O1: thyme) associated with a buried food reward (1/8^th^ of Honey Nut Cheerio). Each trial began with the mouse confined in the central starting cylinder equidistant to the four odour pots. Timing started when the cylinder was lifted, and the mouse was free to explore the arena until it chose to dig in one of the pots. Digging was defined by purposefully moving the shavings with the paws. If an incorrect choice was made and/or prevent multiple digging choice, the mouse was cornered with a grid and the trial was considered terminated. A trial was also terminated if no choice was made within 3 min from the lifting of the starting cylinder and it was recorded as an *omission*. If the animal had two omission trials in a row, digging was reinstated by placing a pot of unscented shavings with a well exposed food reward at the center of the arena. All pots were removed from the maze and rebaited, if necessary.

Once criterion was met in the discrimination phase the mice were moved to the reversal phase. All shavings were replaced with new shavings to prevent discrimination via unintentional cues. O4 (cardamom) was swapped for a novel odour (O5: eucalyptus) and in this phase the rewarded odour was O2 (clove). *Perseverative errors* were choices to dig in the pot with the odour rewarded in the discrimination phase (O1: thyme). *Irrelevant errors* were choices to dig in the pot with the odour that was never rewarded (O3: rosemary). *Novel errors* were choices to dig in the pot with the newly introduced odour (O5: eucalyptus), which was also never rewarded. *Omissions* were trials without a digging choice within 3 min from the start. Total errors are the sum of perseverative, irrelevant, novel and omission errors.

## Supplemental information legends

**Figure S1. Intact presynaptic dopamine D2 receptor function, striatal DA biochemistry and density of DA cell bodies in the eIF4E Tg mice. (a)** Peak concentrations of evoked DA release in the presence of the D2R-like antagonist sulpiride (2 µM) expressed as percentage of the baseline in aCSF. Unpaired, two-tailed *t-*test, t_12_=0.9, ns. N= 9-8 slices/ 3 mice/ genotype. (**b**) Representative Western Blot (WB) images (**c**) and quantification of tyrosine hydroxylase (TH), vesicular monoamine transporter2 (vMAT2) and dopamine reuptake transporter (DAT) in striatal homogenates. Proteins levels were expressed as ratio of DARPP-32 and as a percentage of wild-type controls (WT). TH: Unpaired, two-tailed *t-*test, t_9_=1.6, ns.; vMAT2: Unpaired, two-tailed *t-*test, t_9_=0.1, n.s.; DAT: Unpaired, two-tailed *t-*test, t_9_=1.1, n.s. N=5-6 mice/genotype. (**d**) Quantification of DA and its metabolites DOPAC and HVA in striatal tissues obtained with liquid chromatography tandem mass spectrometry (UHPLC-MS/MS). DA: Unpaired, two-tailed *t-*test, t_10_=0.6, ns.; DOPAC: Unpaired, two-tailed *t-*test, t_10_=0.9; n.s. HVA: Unpaired, two-tailed *t-*test, t_10_=0.3, n.s.; N= 6 mice/ genotype. (**e**) Representative confocal images and (**f**) quantification of the number of midbrain neurons immunolabelled for TH per µm^2^. Unpaired, two-tailed *t-*test, t_30_=0.9, ns. N= 14-18 slices/ 4 mice/ genotype.

**Figure S2. Functional *Cre* expression is confined in striatal DA axons and ACh interneurons in DAT-*Cre* and VGLUT3-*Cre* mice, respectively.** (**a**) Representative confocal images of the striatum of a DAT-*Cre* mouse injected with AAV5-EF1a-DIO-hChR2(H134R)-mCherry-WPRE-HGHpA and stained with antibodies against TH and red fluorescent protein (RFP). DA terminals stained by TH colocalize with RFP labelling mCherry expressed by the AAV5. (**b**) Representative confocal images of the striatum of a VGLUT3-*Cre* mouse injected with AAV5-Ef1a-DIO-EYFP and stained with antibodies against green fluorescent protein (GFP) and choline acetyltransferase (ChAT) expressed by ACh interneurons. (**c**) Representative pie chart illustrating the percentage of YFP positive (YFP^+^) neurons also positive (ChAT^+^) or negative (ChAT^-^) for ChAT.

**Figure S3 Unaltered striatal acetylcholine (ACh) levels and iAChSnFr sensitivity in the eIF4E Tg mice.** Quantification of striatal (**a**) acetylcholine and (**b**) choline in striatal tissues. Unpaired, two-tailed *t-*test, t_43_=0.6, ns. N= 6 mice/genotype. (**c**) Quantification of the changes in fluorescence of the iAChSnFr in striatal slices of wild-type mice following bath application of 10, 25, 50, 100 µM ACh. One-way RM ANOVA F_(1,588, 3,176)_ = 8.2, p=0.05. N= 3 slices/2 mice. (**d**) Quantification of the changes in fluorescence of the biosensor iAChSnFr in striatal slices of wild-type (WT) and eIF4E Tg mice (TG) mice following application of 50 µM of exogenously applied Ach. Unpaired, two-tailed *t-*test, t_18_=0.3, ns. N= 10 slices /2mice /genotype.

