## Supplementary figures and images for "Dysregulated acetylcholine-mediated dopamine neurotransmission in the eIF4E Tg mouse model of autism spectrum disorders"

### Supplementary Figure 1

Supplementary figure 1

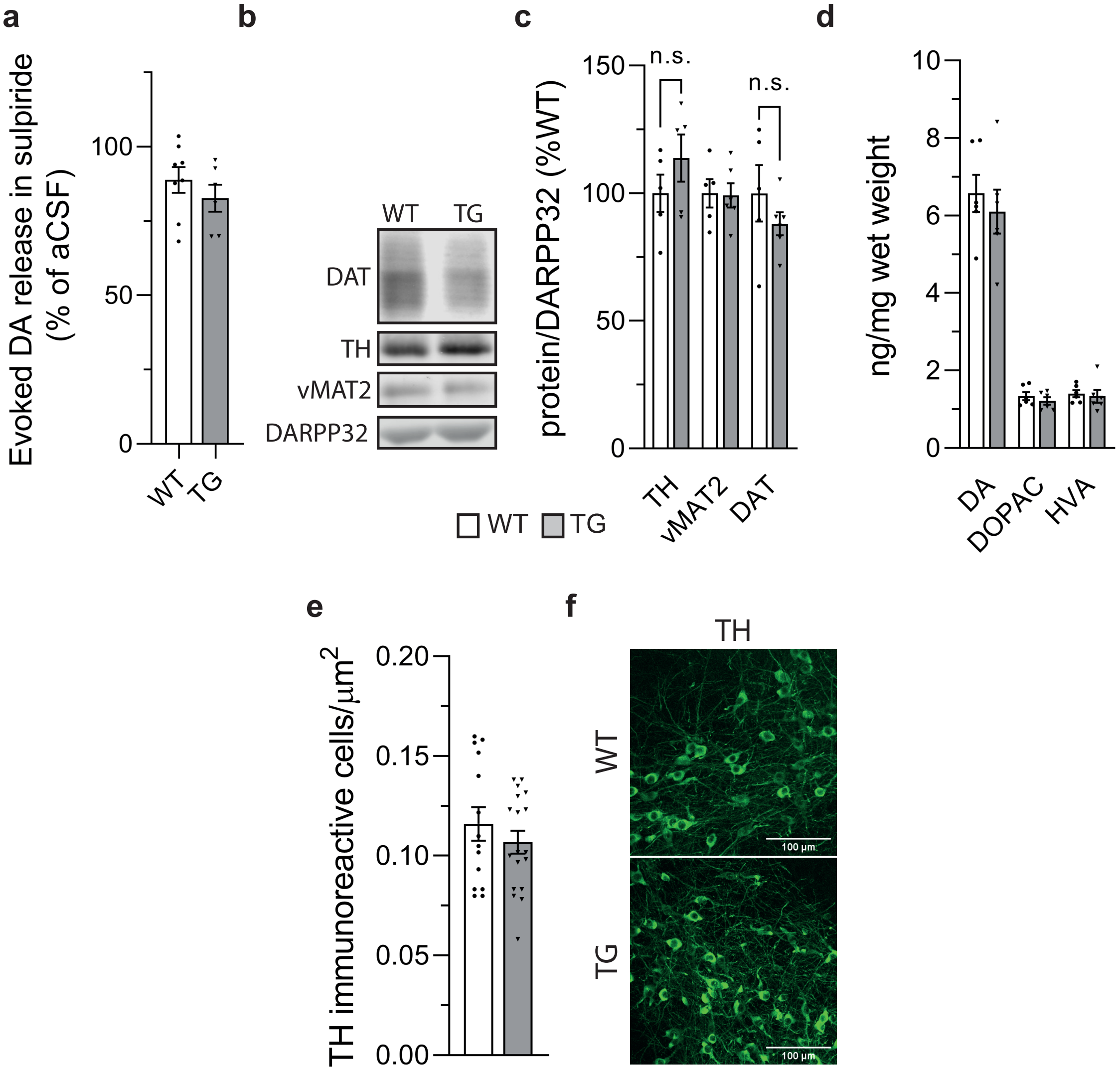

### Supplementary Figure 2

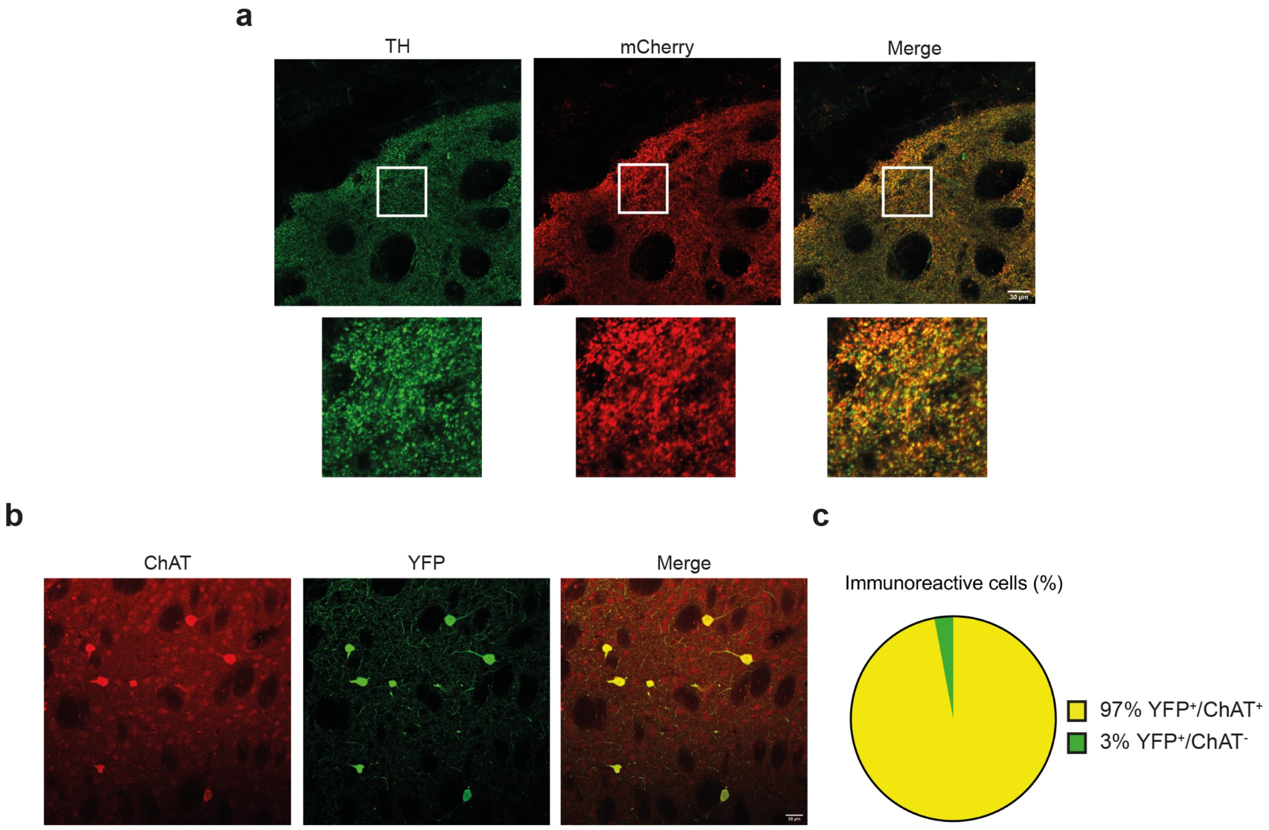

### Supplementary Figure 3

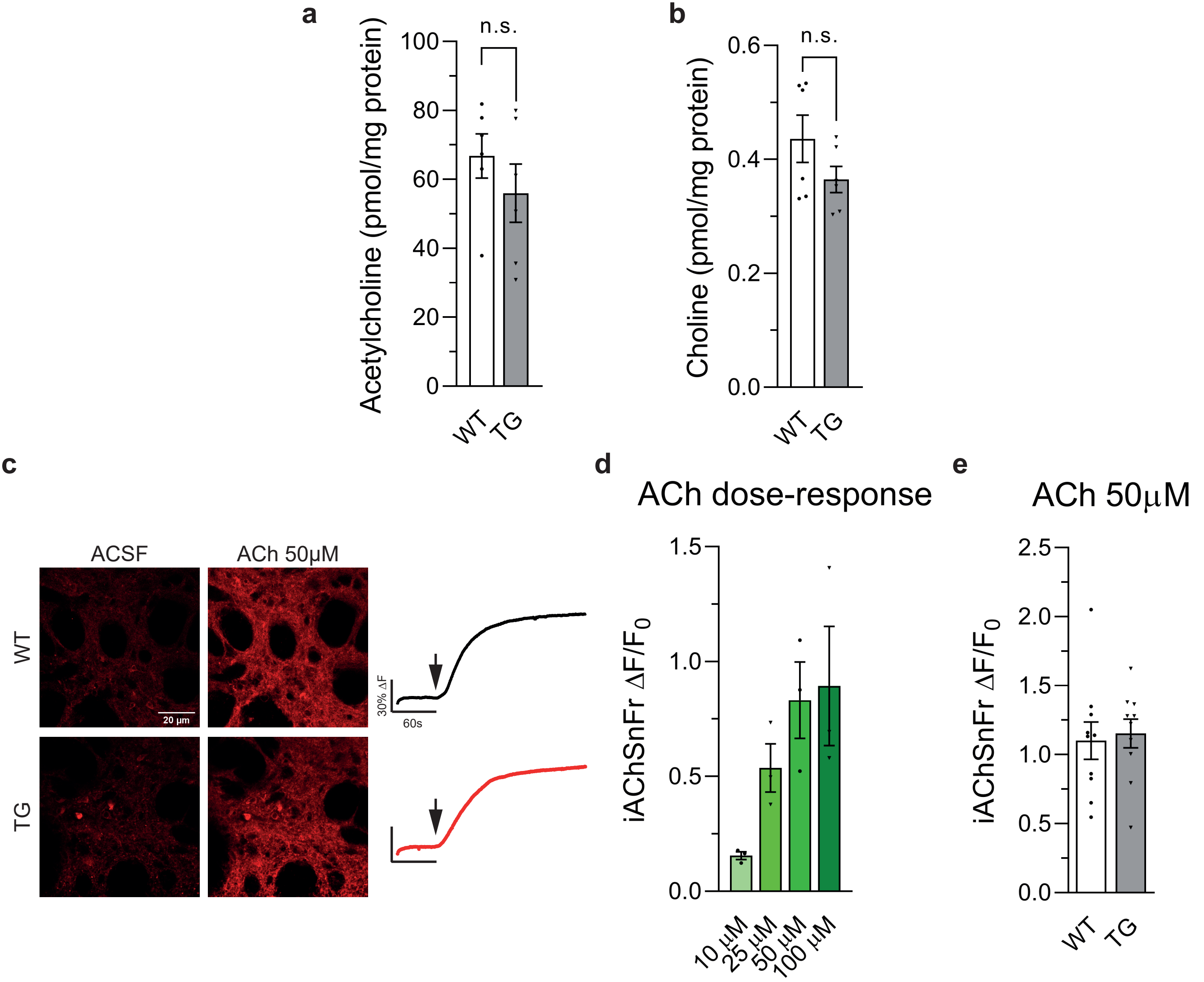
